# Integrated Framework for Multiscale Microvascular Models

**DOI:** 10.64898/2026.04.13.718340

**Authors:** Aref Valipour, Alisyn R. Bourque, Stephen N. Housley

## Abstract

Microvasculature networks mediate nutrient delivery, waste removal, and drug distribution, yet current microfluidic devices fail to capture biological complexity. Here, we introduce an integrative framework to automate generation of bio-informed microvasculature models unifying *in silico* and *in vitro* applications. Our approach leverages a new stochastic growth algorithm governed by fundamental angiogenic principles to generate closed-circuit, fabrication-ready architectures with physiological relevance. We introduce an inverse design strategy that provides a principled mechanism to assign vessel characteristics that satisfy physiological scaling laws. We then present electrical network dynamics, a new algorithm that characterizes network behaviors 100-10,000X faster than CFD, while preserving quantitative predictions. We demonstrate models are fully interchangeable between experimental domains through systematic investigation of vascular topology influence of flow, transport, and cellular behavior. Our platform closes a long-standing gap and provides a generalizable foundation for studying microvascular function in health and disease.

## Introduction

Microvascular networks form the dominant interface for transport and exchange in biological tissues, governing the delivery of oxygen, nutrients, hormones, and immune cells, while simultaneously enabling waste removal and biochemical signaling. Beyond serving as passive conduits, these networks actively shape tissue physiology. Vascular architecture and flow dynamics govern endothelial mechanotransduction(*1*), barrier integrity(*2*), inflammatory signaling(*3*), thrombosis(*3*, *4*), the spatial distribution of drugs(*5*), and metastatic dissemination(*6*). Dysregulation of microvascular structure and/or function is a hallmark of numerous pathologies(*7*, *8*), including cancer(*9*), diabetes(*10*), ischemic disease(*7*), neurodegeneration(*11*), and Alzheimer’s disease(*12*). Consequently, there is strong and growing interest in experimental and computational platforms that can test the microvasculature mechanisms that influence health and pathological cascades.

Progress in this area has traditionally relied on animal models and intravital imaging, which provide physiological relevance but afford limited control over vascular geometry and flow(*13*, *14*). These constraints complicate causal interrogation of how specific architectural features influence function and hinder systematic exploration of fundamental principles. Microfluidic and organ-on-chip technologies have emerged as powerful in-vitro alternatives, enabling precise control over flow dynamics, biochemical cues, and cellular composition. Elegant studies have recreated endothelialized microchannels(*15*, *16*), shear-dependent vascular responses(*17*), and simplified representations of microvascular transport(*16*, *18*). However, most platforms rely on highly idealized geometries(*19*, *20*), single straight channels(*21*), parallel channel arrays(*22*, *23*), isolated bifurcations, or small hand-drawn branching trees(*24*, *25*) integrating only divergences that capture fragments of the topological complexity inherent in biology.

A key limitation is the implicit assumption that flow through each vessel is determined locally by its diameter and length. In vivo, microvessels exist within densely interconnected networks in which their topology including loop density, anastomoses, and hierarchical connectivity can generate heterogeneous shear stresses, preferential flow routes, and nonlinear redistribution following occlusion(*26*). These emergent, network-level behaviors are central to physiological and pathological phenomena, yet they are inaccessible with current methods(*27*). At the same time, a persistent gap separates computational and experimental approaches. In-silico studies often employ idealized networks optimized for numerical efficiency but are incompatible with microfabrication. Conversely, experimentally fabricated devices are frequently simplified in their network complexity and their minimal cross-sectional area to accommodate lithography or imaging. These compromises limit their utility for rigorous simulation and the biologic scale at which inferences can be derived(*18*, *25*). As a result, direct comparisons between computational predictions and experimental measurements are lacking. When discrepancies arise, it is difficult to disentangle physical effects, e.g. non-Newtonian rheology(*28*, *29*), wall compliance(*30*, *31*), or cell–fluid interactions(*32*, *33*) from artifacts introduced by geometric mismatch. This disconnect has slowed validation and impeded the development of predictive, generalizable models of microvascular transport.

Existing vascular network generation algorithms underscore this challenge. Methods based on constrained constructive optimization(*34*) and related frameworks successfully reproduce the acyclic, diverging topologies seen in arterial and venous trees but fail to reproduce capillary beds, where closed loops (converging and diverging) and anastomotic connections are ubiquitous and essential to function(*24*, *35–37*). Moreover, detailed computational fluid dynamics simulations of complex networks are computationally expensive, often requiring hours to days for networks of realistic size. This cost precludes large parametric studies, rapid design iteration, and systematic exploration of how architectural features shape function. Consequently, there is no framework for generating closed-circuit microvascular networks that are simultaneously biologically informed, computationally analyzable, and directly fabricable.

Here, we introduce an integrative framework for generating bio-informed microvasculature models that unifies *in silico* and *in vitro* applications (Fig. 1). Our approach leverages a stochastic growth algorithm governed by fundamental angiogenic principles to generate closed-circuit, fabrication-ready architectures with physiological relevance, a first for the field. We introduce inverse design strategy to enforce Murray’s law in closed-circuit networks, providing a principled mechanism to automate assignment of vessel characteristics that satisfy physiological scaling laws within complex, circular dependencies. We present electrical network dynamics (END), a new algorithmic framework that characterizes network behaviors orders of magnitude faster than traditional CFD, while preserving quantitative predictions. We demonstrate that models are fully interchangeable between computational and experimental domains through systematic, reproducible investigation of how vascular topology governs flow, transport, and cellular behavior. Collectively, our platform closes a long-standing gap between theoretical prediction and empirical studies by providing a generalizable foundation for studying microvascular function in health and disease.

**Fig. 1:**
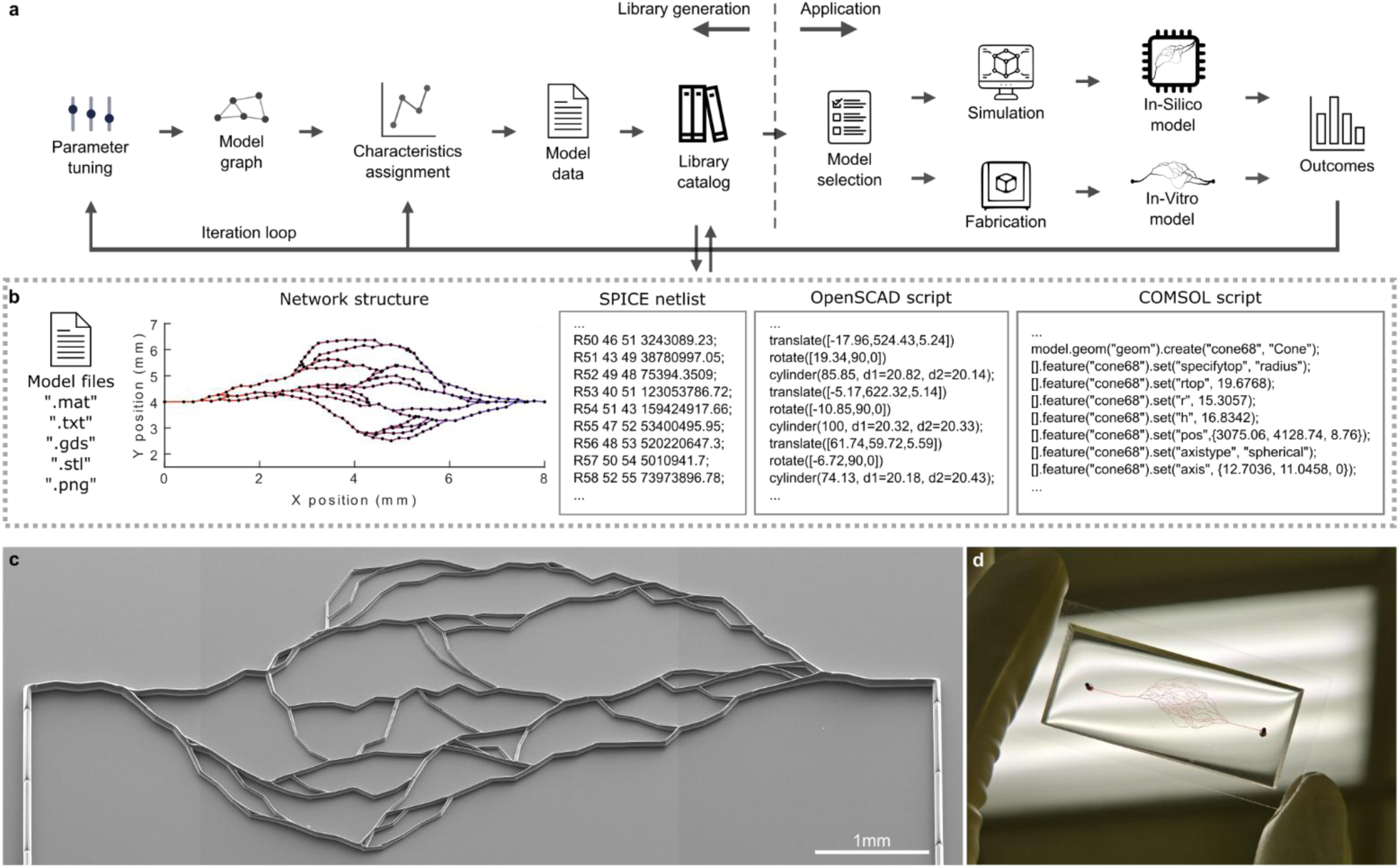
Integrated workflow of the microvasculature platform bridges computational design, in vitro, and in silico experiments. (**a**) Complete pipeline from algorithmic generation to application, divided into library generation (left) and application (right) phases. Library generation: Parametric tuning of control variables drives a stochastic growth algorithm generating network graphs with node coordinates, connectivity matrices, and segment geometries. Automated translation pipelines convert each graph into: (1) CAD files (STL, GDSII) for fabrication, (2) geometries for CFD, and (3) SPICE netlists for END. CFD characterizes hemodynamics (velocity, pressure, and shear stress), while END solves network-wide flow distribution orders of magnitude faster than CFD. Each model entry in the library catalog contains generation parameters, topology data, CAD files, SPICE netlist, and basic characteristics. Application: Researchers select models based on architectural requirements (e.g. vessel density, tortuosity, diameter distribution). Selected models support two pathways enabling cross-validation: In-Silico models for computational studies using CFD (detailed fluid dynamics, particle trajectories) or END (rapid parametric screening), and In-Vitro models for physical fabrication via soft lithography (GDSII) or two-photon polymerization (STL) to create functional microfluidic devices. Identical geometries in simulations and fabricated devices enable direct cross-validation, eliminating geometric variability as a confounding factor. (**b**) Each catalog entry provides: (i) rendering views of the network architecture, (ii) quantitative metrics including vessel density, branching statistics, connectivity matrices, and tortuosity measurements, and (iii) implementation-ready files including SPICE netlists for electrical analog simulation, OpenSCAD scripts for rendering 3D fabrication files, GDSII masks for soft lithography, and COMSOL geometries for CFD analysis. Scripts are shortened for visual representation. (**c**) A tiled electron micrograph (SEM image) of a sample fabricated vascular structure. (**d**) A sample fabricated microfluidic vascular chip with colored dies infused for contrast.

## Results

### Stochastic growth algorithm generates biologically faithful microvascular networks

We developed a stochastic growth–based algorithm that generates microvascular networks constrained by core biological principles of angiogenesis while remaining fully parametric and computationally tractable. Unlike bifurcating trees, microvascular beds are characterized by dense anastomoses, enabling redundant perfusion and adaptive redistribution of flow. Existing synthetic models do not capture this feature.

Our framework integrates eight fundamental control variables governing network geometry and topology, including branching probability, step size, anastomosis distance, and diameter scaling (Fig. 2a)(*38*, *39*). Our approach explicitly incorporates both diverging and converging vessel formation during network growth, yielding architectures that are topologically distinct from conventional tree-based designs. Networks are grown from predefined inlets via probabilistic advancement of vessel tips, which may extend, branch, or anastomose depending on local geometric context. Anastomoses arise either through direct intersection with existing vessels or proximity-based connections, while a loop-size constraint suppresses nonphysiological microloops. This growth process naturally produces heterogeneous capillary-scale architectures with variable segment lengths, branching patterns, and loop densities that closely resemble native microvasculature.

**Fig. 2:**
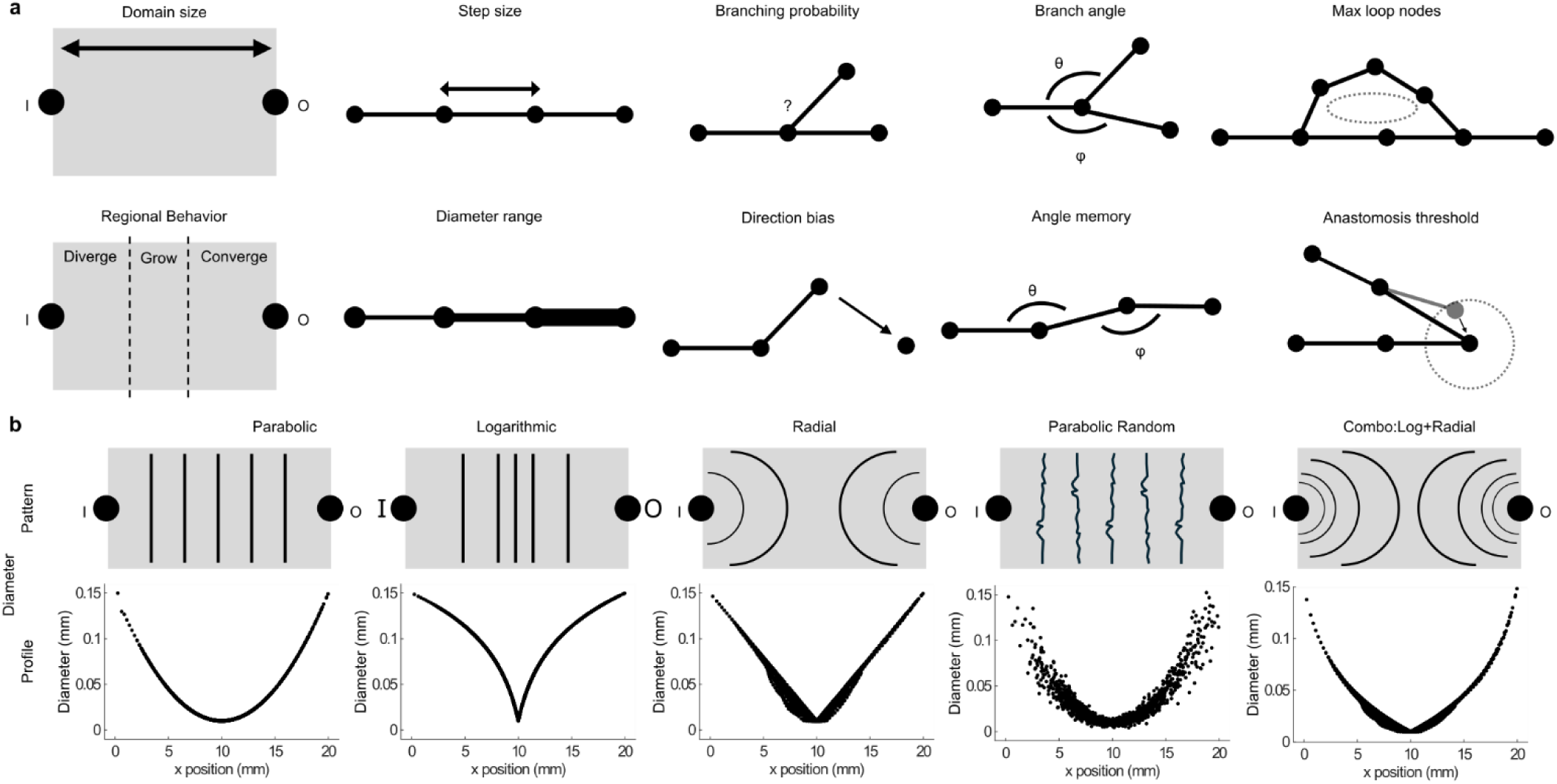
Parametric control framework enables functional tuning of network characteristics. (**a**) Two domain specifying and eight fundamental control parameters govern network topology and geometry: domain size (tissue region extent), step size (segment length), branching probability (affecting capillary density), branch angle (directional range), maximum loop nodes (loop size constraint), regional behavior (spatial variation in growth rules dividing domain into divergent, growth, and convergent zones with distinct branching and directional properties), vessel diameter range (size distribution bounds), direction bias (preferential growth toward outlet), angle memory (directional persistence from parent vessel), and anastomosis threshold (proximity distance for branch merging). Regional behavior modulates growth characteristics as vessels advance from inlet (I) to outlet (O), with early regions exhibiting high directional randomness, middle regions showing increased branching probability and wider angular variation to maximize capillary density, and late regions displaying reduced branching with narrower angles for convergence toward outlets. (**b**) Control variables can be expressed as mathematical functions of spatial position rather than static constants, enabling precise spatial patterning. Five representative diameter assignment functions demonstrate this capability: parabolic profiles create high contrast with maximum diameters at horizontal domain edges transitioning to minimum at center; logarithmic functions produce gradual diameter gradients; radial functions generate concentric size patterns mimicking arteriolar-to-capillary transitions; parabolic with 20% random noise introduces realistic biological variability while maintaining overall spatial trends; combined radial-logarithmic functions create complex hierarchical organizations. Top row shows simple illustrative schematic patterns with inlet (I) and outlet (O) positions. Bottom row displays resulting diameter profiles along the x-axis (normalized position 0-20 mm), demonstrating how mathematical functions translate into vessel size distributions across the network domain.

Crucially, parameters can be defined as spatial functions rather than constants, enabling hierarchical organization within a single network (Fig. 2b). By modulating controlled randomness at multiple hierarchical levels, primary stochasticity for trajectory and secondary randomness for local irregularities, we balance biological variability with reproducibility. For example, modulating branching probability generated regions of high capillary density flanked by larger upstream and downstream vessels. Networks generated with identical parameter sets but different random seeds exhibit diverse topologies while preserving consistent statistical properties, including segment count, loop density, and path length (Fig. 3).

**Fig. 3:**
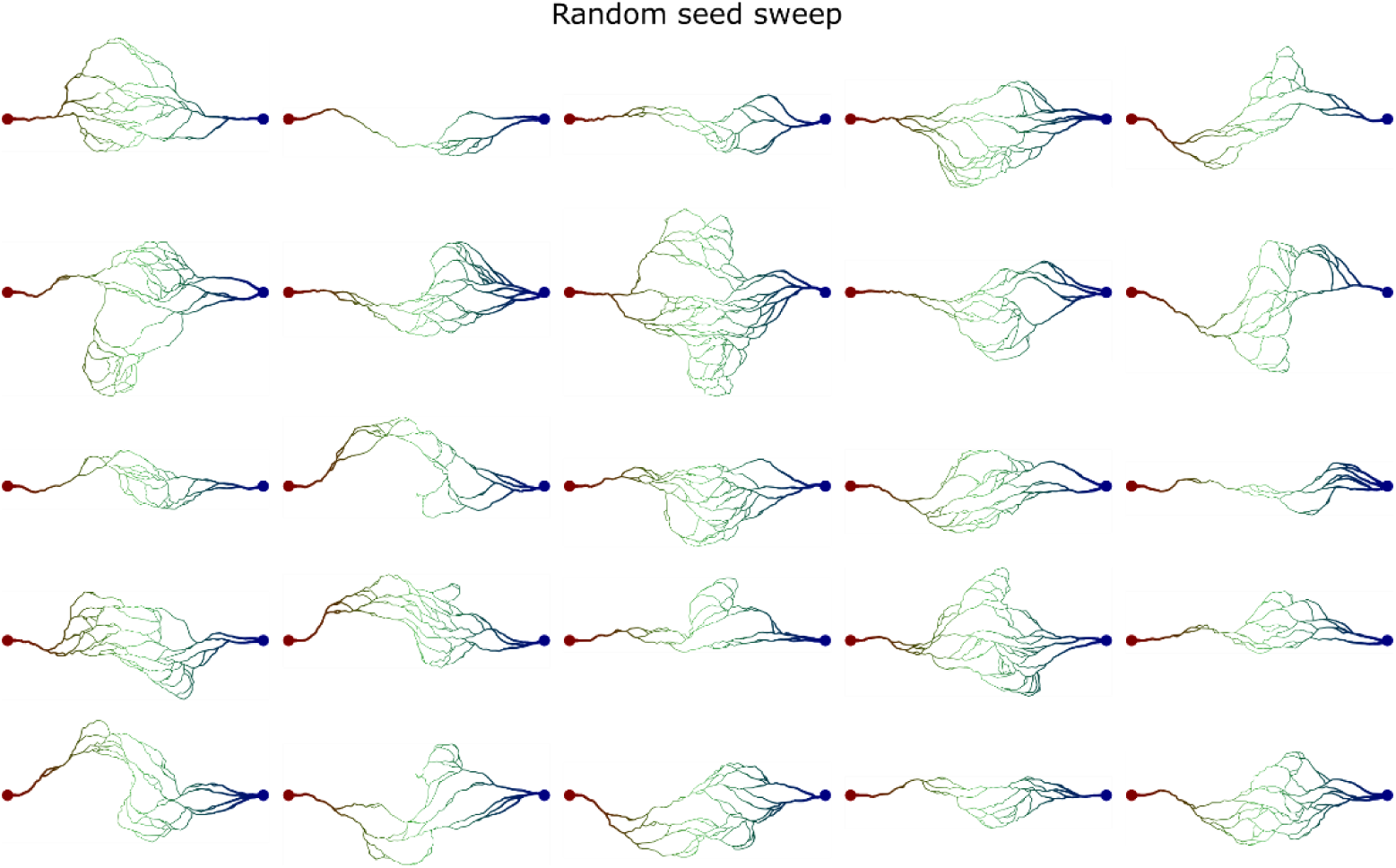
Models generated with a sweep on random seed. The variations suggest the inherent stochastic nature of the algorithm translating into the difference in models generated from the same parameters. Random seed range is [101:125]. Model parameters are: domain size: 15 mm, step size: 250 µm, branch probability: 0.2, branch memory strength: 0.35, outlet attraction start: 0.35, outlet influence maximum: 0.6, middle region start: 0.2, middle region end: 0.6. Colors: red represents proximity to arteriolar inlet vessels and blue represents proximity to venular outlet vessels.

### Unified translation pipelines link topology, simulation, and fabrication

A key advance of our platform is the ability to translate identical large-scale network graphs into multiple downstream representations without loss of geometric or topological fidelity. We developed four automated pipelines converting network graph representations into: (1) 3D CAD models for volumetric high-resolution printing; (2) GDSII photomasks for soft lithography; (3) SPICE netlists for END analysis; and (4) COMSOL geometries for Multiphysics simulations. For 3D applications, the pipeline generates OpenSCAD or COMSOL scripts that construct truncated cones with spherical junctions to ensure watertight geometry. Conversely, for planar applications, the GDSII pipeline creates photomasks compatible with conventional lithography, automatically adding port structures for macro-to-micro interfacing.

Because all outputs derive from the same underlying graph, computational predictions and experimental devices are geometrically identical. This interoperability allows the same network topology to be analyzed via END, verified with 3D CFD, and fabricated using either two-photon polymerization or soft lithography, ensuring that discrepancies are attributable to physical phenomena rather than geometric inconsistencies. Our modular architecture further enables future extensions without disrupting existing workflows.

### Systematic exploration of architecture–topology relationships reveals biological diversity

We then explored the range of architecture–topology relationships by systematically varying individual control parameters while holding others constant. These systematic parameter sweeps revealed clear relationships between design parameters and emergent network properties. Increasing domain size scaled network complexity proportionally, transitioning from isolated capillary beds to tissue-scale microvascular meshes. Varying branching probability controlled a continuum from sparse, tree-like networks to dense, highly interconnected architectures, with intermediate values producing morphologies most consistent with physiological capillary beds. Step size regulated spatial resolution and segment length distributions, whereas advancing angular modulated vessel tortuosity. Collectively, these sweeps define a structured design space linking intuitive parameters to quantitative network metrics. Each generated network is cataloged with its full parameter set and associated topological statistics, forming a reusable library of microvascular topologies. This resource enables investigators to select or generate networks tailored to specific experimental or computational requirements without manual redesign.

### Inverse design strategy enforces Murray’s law in closed-circuit networks

A major challenge in vascular engineering is assigning vessel diameters that satisfy Murray’s Law(*40*) in closed-circuit networks, where circular dependencies make traditional parent-child calculations impossible. We resolved this by developing an inverse design approach that exploits the mathematical isomorphism between fluidic networks and electrical circuits governed by Kirchhoff’s Current Law (KCL) (Fig. 4a-c).

**Fig. 4:**
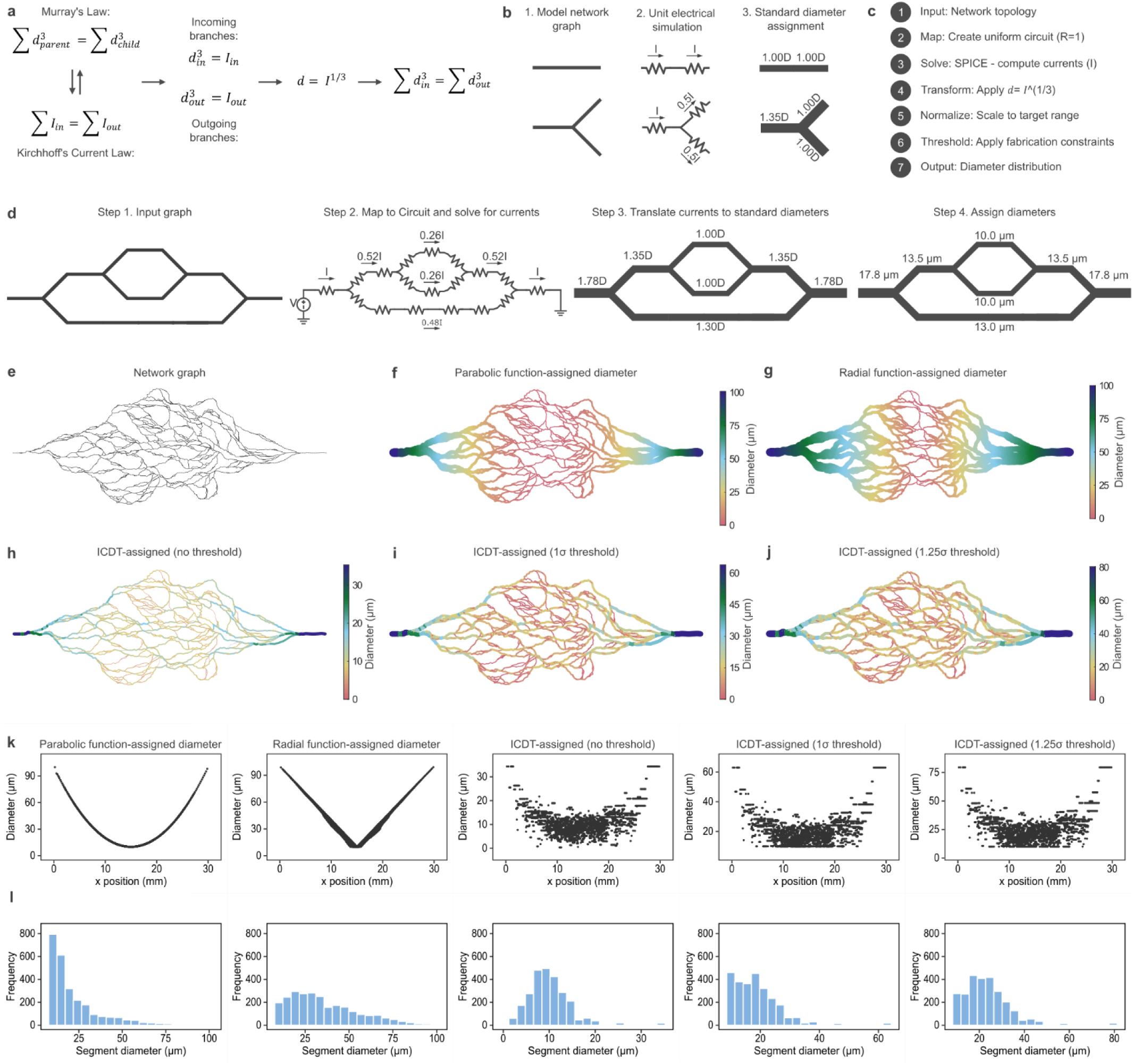
Principal mechanism for Inverse Current-to-Diameter Transformation (ICDT). (**a**) Theoretical foundation: Murray’s Law relates parent and child vessel diameters at bifurcations (top), while Kirchhoff’s Current Law ensures flow conservation (bottom), enabling electrical circuit analogy for fluid networks. (**b**) In a straight channel, all the resistors (segments) will have the same current (flow rate) and end up with same diameter. In Y-junctions (or higher degree connections), the currents (flow rates) follow KCL, therefore one segment (parent segment) will have a larger diameter than others connected to the same node (daughter segments). (**c**) Seven-step computational pipeline of ICDT. (**d**) Workflow schematic showing sequential processing steps of ICDT. The output of ICDT is a set of standardized diameter ratios, scalable to the desirable diameter ranges. (**e**) Example microvascular network graph showing complex branching topology. (**f-g**) Function-based diameter assignment methods: parabolic function creates symmetric diameter gradients from boundaries, while radial function assigns diameters based on distance from network inlet/outlet. (**h-j**) ICDT-assigned diameters with varying fabrication constraint thresholds: (**h**) no threshold maintains full diameter range, (**i**) 1σ threshold, and (**j**) 1.25σ threshold remove smaller vessels below fabrication limits. Quantitative comparison of diameter assignment methods showing (**k**) spatial diameter profiles and (**l**) resulting diameter distributions. Parabolic and radial methods produce deterministic gradients, while ICDT generates biologically realistic diameter heterogeneity.

We first mapped network topology to an equivalent electrical circuit with uniform resistance, ensuring that current distribution depends solely on connectivity. Steady-state branch currents are then calculated via SPICE analysis. These calculated branch currents are then transformed into vessel diameters using Murray’s power-law relationships (i.e., d ∝ I^1/3^ for capillaries), and the resulting distribution is normalized to a user-defined scale (Fig. 4d). This approach guarantees global consistency at all junctions, including within closed circuits where direct application of Murray’s law would otherwise require burdensome computations.

Our automated method generates internally consistent diameter hierarchies for arbitrary network topologies in seconds, representing, to our knowledge, the first scalable solution for enforcing Murray’s law in closed-circuit microvascular networks. We contrasted this biological approach with a function-based assignment strategy (e.g., parabolic or radial profiles), which offers higher parametric contrast useful for studying size-dependent phenomena like cell margination. While function-based assignments could generate wide diameter ranges (1–100-fold contrast for a 1000-segment network), our inverse design produces the moderate, smooth transitions (3–6 fold for a 1000-segment network) characteristic of physiological capillary beds (Fig. 4e-l).

### Electrical network dynamics (END) enables rapid network-level screening

We further leveraged fluidic–electric isomorphism (Fig. 5a) to develop electrical network dynamics (END), a new framework for large-scale network characterization of fluid properties without the computational cost of full 3D computational fluid dynamics (CFD). In END, each vessel segment is represented as a resistor whose value is derived from modified Hagen–Poiseuille relationships, while circuit connectivity directly maps to network topology (Fig. 5b-c). Solving the resulting circuits via SPICE (Fig. 5d) yields pressure and flow distributions in milliseconds to seconds, orders of magnitude faster than equivalent CFD simulations. This also allows us to assign a single representative resistance value to any given microvascular network (Fig. 5e). END predictions closely match CFD results for relative flow distributions, accurately identifying high- and low-flow pathways across complex networks confirmed by close match with in vitro particle velocimetry. END’s efficiency enables the large-scale (1000s) screening of models and real-time optimization of network architectures that would be computationally prohibitive using CFD alone.

**Fig. 5:**
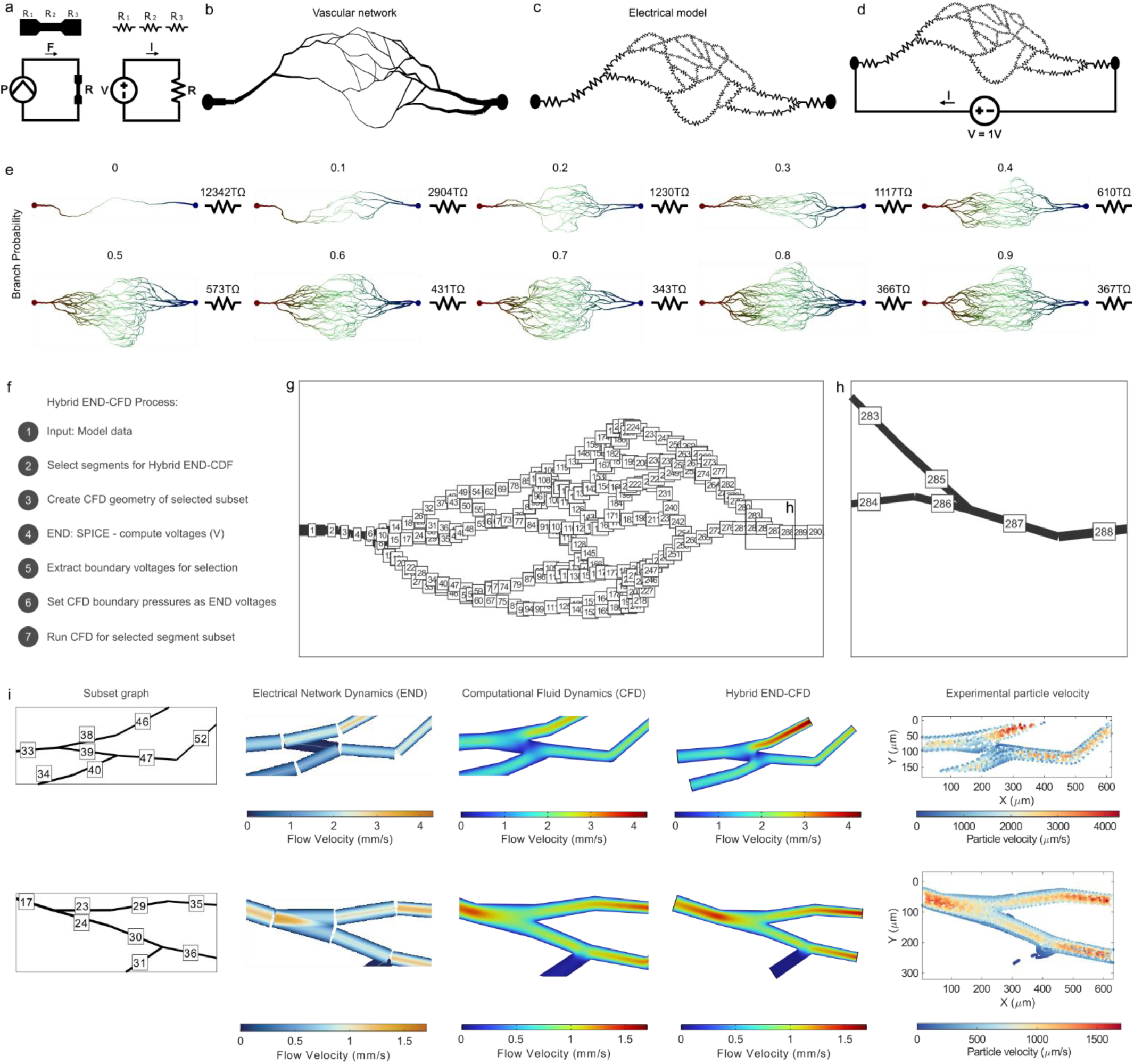
Electrical Network Dynamics (END) enables rapid simulation and hybrid END-CFD framework. (**a**) Electrical circuit analogy for vascular networks: Hydraulic resistance of each vessel segment maps directly to electrical resistance, where pressure difference equals voltage and flow rate equals current. This enables rapid network-scale flow computation using established circuit simulation tools. (**b**) Generated network graph containing node coordinates, connectivity matrix, and segment geometries. (**c**) Electrical network model: each vessel segment becomes a resistor, converting fluid network into solvable electrical circuit. (**d**) Standardized test circuit: 1V DC source drives current through the network. Equivalent network resistance calculated as R_eq_ = 1V / I_Source_, providing single scalar metric for quantitative architecture comparison. (**e**) Equivalent resistance for networks spanning branching probabilities 0-0.9. Sparse networks (low branching) exhibit high resistance due to limited parallel pathways; dense networks (high branching) show reduced resistance as anastomotic loops create multiple flow paths. (**f**) Hybrid END-CFD workflow. (**g-h**) In hybrid END-CFD, a subset (nodes 283-288) of the complete network topology is selected for CFD simulation. (**i**) Cross-validation of simulation methods: Two representative subset of segments analyzed across three simulation approaches and the experimental particle velocimetry as the reference (ground truth). Subset graph shows the selected subgraph. END simulation provides rapid flow velocity estimates from electrical analog. CFD simulation delivers high-resolution velocity fields. Hybrid END-CFD demonstrates seamless integration where CFD subset receives boundary conditions from global END solution, reducing computational cost. Agreement across all methods validates the electrical analog approach for rapid screening while demonstrating CFD’s utility for detailed mechanistic studies. The hybrid approach optimally balances computational efficiency with spatial resolution, enabling network-scale simulations with region-specific high-fidelity analysis. All color bars in a row are set to have the same bounding range for better colormap comparison. (Bottom row represents the same subgraph as fig. 5 for comparison)

### Hybrid END–CFD approach resolves local phenomena within tissue-scale networks

To address the computational intractability of full CFD simulations in large networks (Fig. 6), we developed a hybrid strategy that combines END with localized CFD (Fig. 5f). END analysis of the complete network (Fig. 5g) provides nodal pressure boundary conditions, which are then applied to selected subgraphs (Fig. 5h) representing regions of interest. High-resolution CFD is then performed only on these subdomains, reducing computational burden by up to three orders of magnitude while preserving physiologically accurate boundary conditions. Comparisons with full-network CFD and END demonstrate that this hybrid approach accurately reproduces local pressure and velocity fields, enabling detailed investigation of shear-dependent and junction-level phenomena within the context of realistic tissue-scale networks (Fig. 5i).

**Fig. 6:**
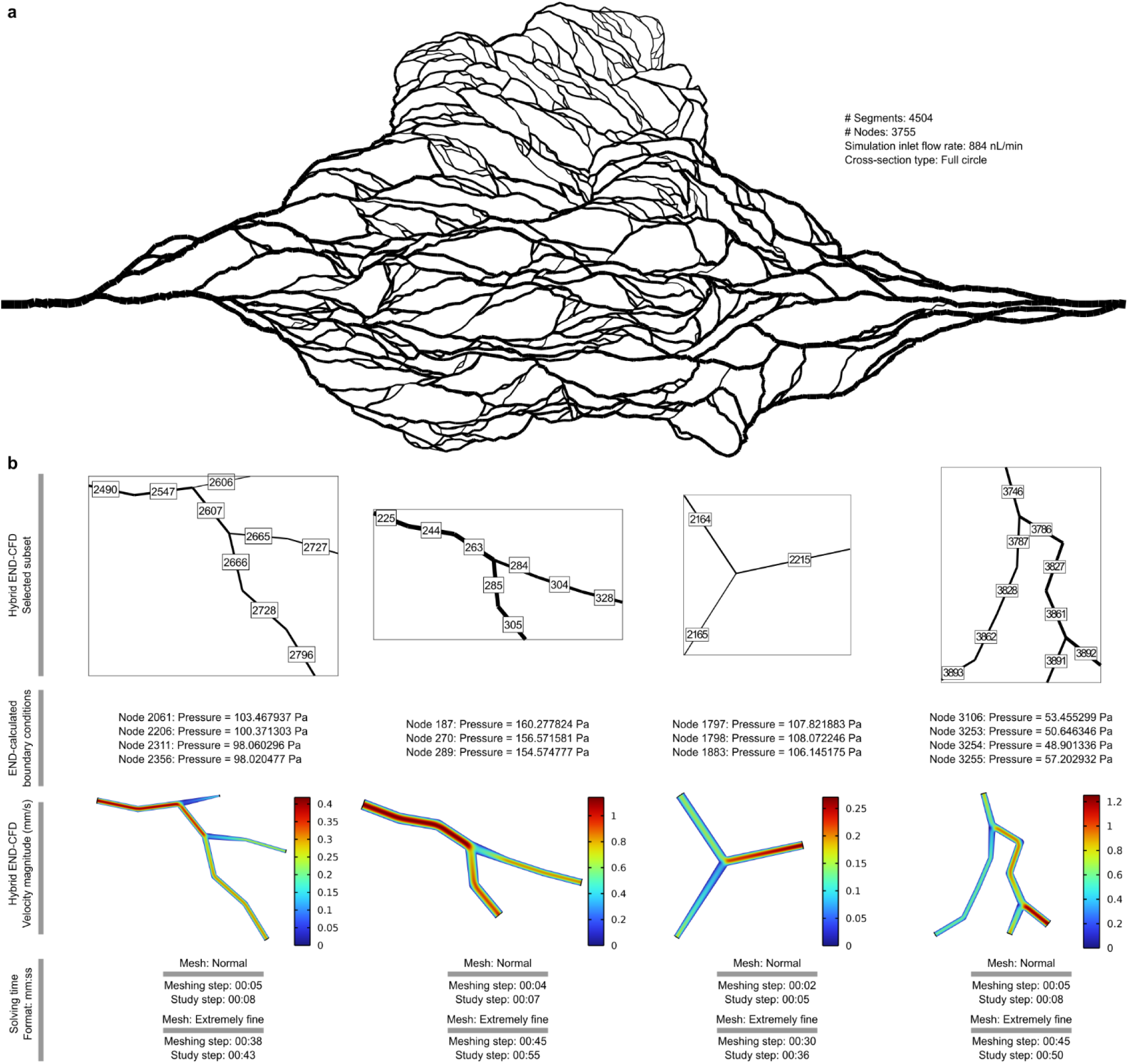
Hybrid END-CFD enables high-resolution in-silico study of large vascular networks. (**a**) Larger vascular networks are costly and often unfeasible to simulate using CFD alone. A representative large microvascular network is shown that consists of 4504 segments and 3755 nodes. (**b**) Selected subsets are simulated with our Hybrid END-CFD approach. END simulation calculates the boundary pressures for CFD. CFD simulations are done using “Normal” and “Extremely fine” mesh qualities. The time for mesh generation or simulation of the study for above-illustrated sub-graphs were both under one minute, confirming fast turnaround of the Hybrid END-CFD pipeline.

### Network topology, not local geometry, governs flow heterogeneity

We then utilized our new unified framework to explore a key question in microvascular topology-function relationships. By comparing simple tree-like architectures with complex anastomotic networks composed of geometrically identical segments, our analysis demonstrates that flow behavior in microvascular networks depend on network-level architecture rather than isolated, local segment geometry (Fig. 7a-f). Even when all segments share identical dimensions, differences in downstream resistance produce unequal flow distributions, an effect amplified by the presence of loop dependencies confirmed by END and in vitro observations. Local geometry then translates this to local characteristics. Flow velocity is set at segment level based on the flow rate set at the network level. CFD simulations confirm these predictions, establishing that local flow conditions are dictated by global network architecture rather than by isolated channel geometry alone (Fig. 7g). This result underscores a critical limitation of conventional microfluidic designs based on isolated bifurcations or parallel channels and highlights the necessity of network-level models to capture emergent hemodynamic phenomena intrinsic to biological microvasculature.

**Fig. 7:**
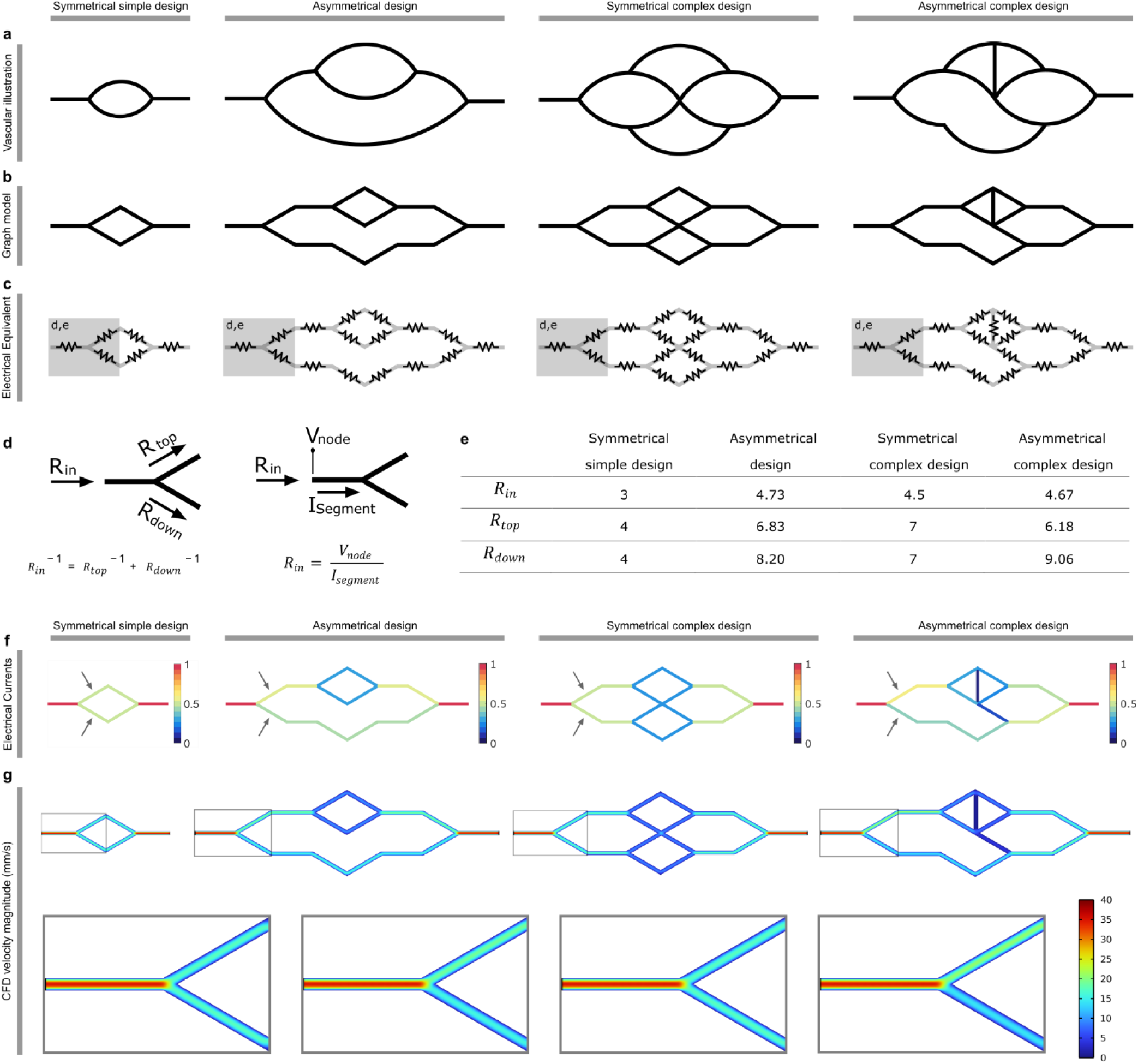
Network topology, not local geometry, governs flow heterogeneity. (**a**) Vascular design archetypes: Four representative network configurations spanning symmetrical simple (single bifurcation, representing current Y-channel vessel models), asymmetrical (unequal branching), symmetrical complex (nested parallel bifurcations, representing current more advanced vessel models), and asymmetrical complex (combined branching and structural asymmetry) designs. These represent fundamental building blocks of microvascular model architecture. All segments are assumed to have the same dimensions. (**b**) Corresponding graphs represent topological connectivity and branching structure for each design archetype. (**c**) Electrical equivalent circuits with resistor networks representing hydraulic resistance of each vessel segment. Shaded boxes indicate bifurcation regions (d,e) analyzed in detail. (**d**) Input resistance calculation methodology: At each bifurcation node, parallel branches combine via reciprocal sum, then add in series with upstream resistance. Alternative formulation using voltage drop across segments enables direct computation from SPICE simulations. (**e**) Quantitative comparison of resistance values across architectures: Symmetrical designs show equal branch resistances, while asymmetrical designs exhibit significant imbalance. Network topology determines the resistance seen at each node. (**f**) Standardized electrical current distribution shows flow partitioning at bifurcations. Color maps reveal preferential flow patterns: symmetrical designs split current equally (both branches ∼0.5), while asymmetrical designs create dominant pathways (low resistance paths have higher flowrate). (**g**) CFD validation of END predictions: Top row shows velocity magnitude distributions in network geometries. Bottom row demonstrates the difference in velocity field formed in identical inlet channels when there is a difference in flow rates caused by difference in Rin each channel sees. CFD results confirm END-predicted flow partitioning, vessels with lower resistance exhibit proportionally higher flow rates.

### Fabrication-aware design links cross-sectional geometry to network resistance

Next, we quantified how fabrication-constrained cross-sectional geometries influence dynamic behaviors. Using CFD and hydraulic radius-based mappings, we compared circular, rectangular, and partially circular cross-sections across a range of network architectures. We found that absolute resistance depended strongly on cross-sectional geometry and diameter assignment strategy. Crucially, rectangular cross sections, the most commonly utilized in microfabrication, exhibit lower resistance than circular channels of equivalent hydraulic diameter and have higher fractions of stagnant flow owing to their angular geometry. These results provide a practical and quantitative guide for selecting fabrication methods, balancing the experiment specific need for physiological circular cross-sections against the scalability of planar lithography.

### Physical Realization of Microvascular Digital Twins

To empirically validate the predictive capacity of our framework, we fabricated microvascular devices directly from identical network graphs using two complementary microfabrication strategies: two-photon polymerization (2PP) for fully three-dimensional architectures and soft lithography for planar devices. In both cases, fabrication inputs were generated automatically from the same microvascular network representations used for END and CFD simulations, ensuring that experimental observations could be directly cross-referenced with “digital twin” simulations.

For applications requiring complete geometric fidelity, we employed a MATLAB-to-OpenSCAD pipeline to generate 3D CAD models for 2PP. This enabled fabrication of freestanding, microvascular networks with optimal for high resolution microscopy (partially circular) cross-sections, with all the vascular segments laid on the same plane, and continuous diameter variation throughout the network. This maskless lithography technique produced devices that preserved the precise topology and geometric hierarchies defined by framework, reproducing key features of native capillary beds. In parallel, soft lithography is used to fabricate planar microfluidic networks that preserve identical in-plane topology, segment lengths, and diameter hierarchies, enabling higher-throughput experiments. Despite differences in dimensionality and cross-sectional geometry, both fabrication approaches faithfully reproduced the designed network topology and relative diameter distributions(Fig. 8).

**Fig. 8:**
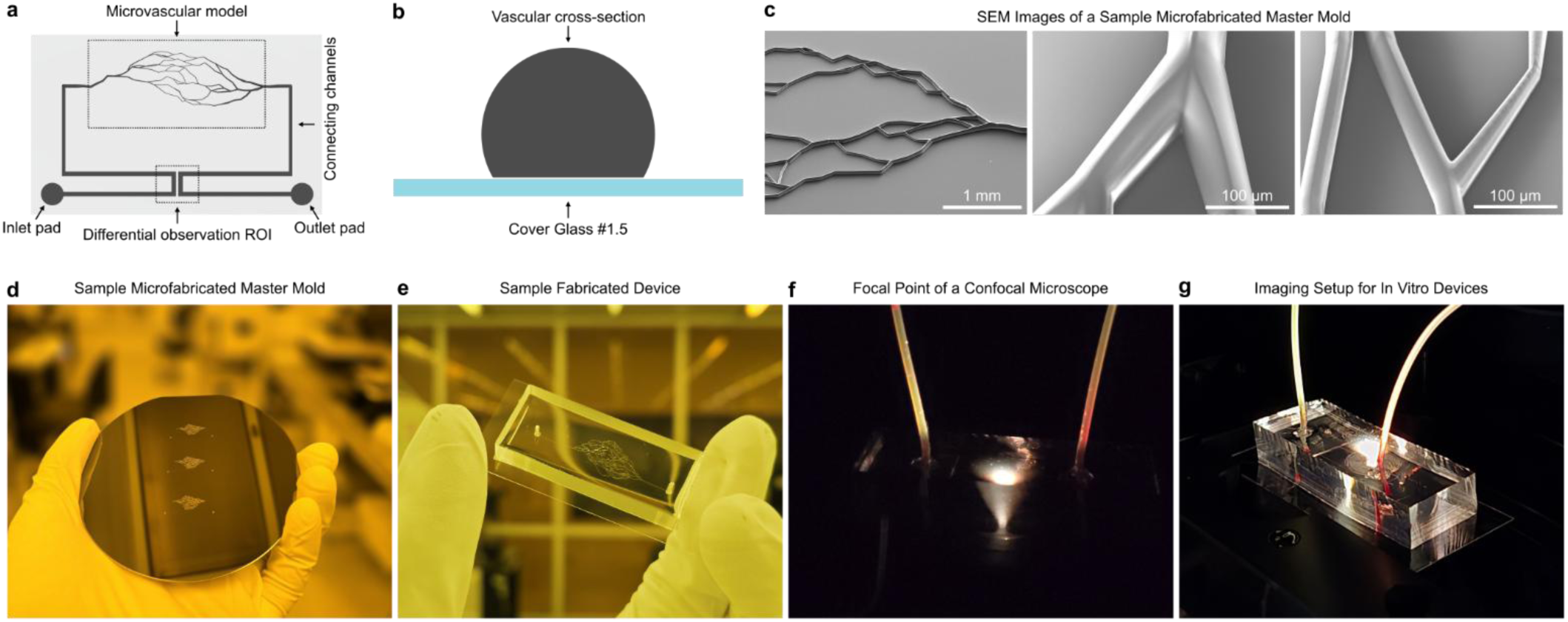
Design and fabrication of in vitro microfluidic devices. (**a**) Helper parts are added to vascular models to facilitate microfluidics utility, including connecting channel, inlet and outlet pads, and dedicated observation region. (**b**) 75% cut cross-section, all laid on the same plane, was selected to allow for high-resolution confocal microscopy within the channels. (**c**) SEM images of microfabricated master mold. (**d**) A sample microfabricated master mold. (**e**) A sample fabricated microfluidic device. (**f**) Focal point of the confocal microscope is at the same height as all segments. The channels are designed laid on the same plain (opposed to center aligned). (**g**) Imaging setup for in vitro experiments.

### Cross-platform validation establishes predictive correspondence from design to experiment

We first cross-referenced device resistance calculated with END, CFD, and measured in vitro. The resistance observed matched closely across in silico and in vitro mediums. At a minimum, these resistance values assist in rapid calculation of pressure-flow required to create intended flow profile within the models.

We then assessed the hydrodynamic accuracy of fabricated networks by perfusing them with fluorescent microparticles to map velocity fields, serving as a physical proxy for the computational flow predictions. Under steady perfusion conditions, experimental particle trajectories were captured via high-speed (∼30-100fps) confocal microscopy, converted into spatially resolved velocity maps, then compared against velocities derived from END and CFD simulations.

Empirical flow fields demonstrated a quantitative match with in silico flow predictions. High-flow and low-flow pathways identified computationally were consistently observed experimentally, and segment-wise velocities closely aligned with model predictions (Fig. 9). These data demonstrate that the END and CFD simulations accurately predict behavior in physical devices and validates our fundamental assumption that network architecture, rather than isolated segment geometry, determines local hemodynamic conditions.

**Fig. 9:**
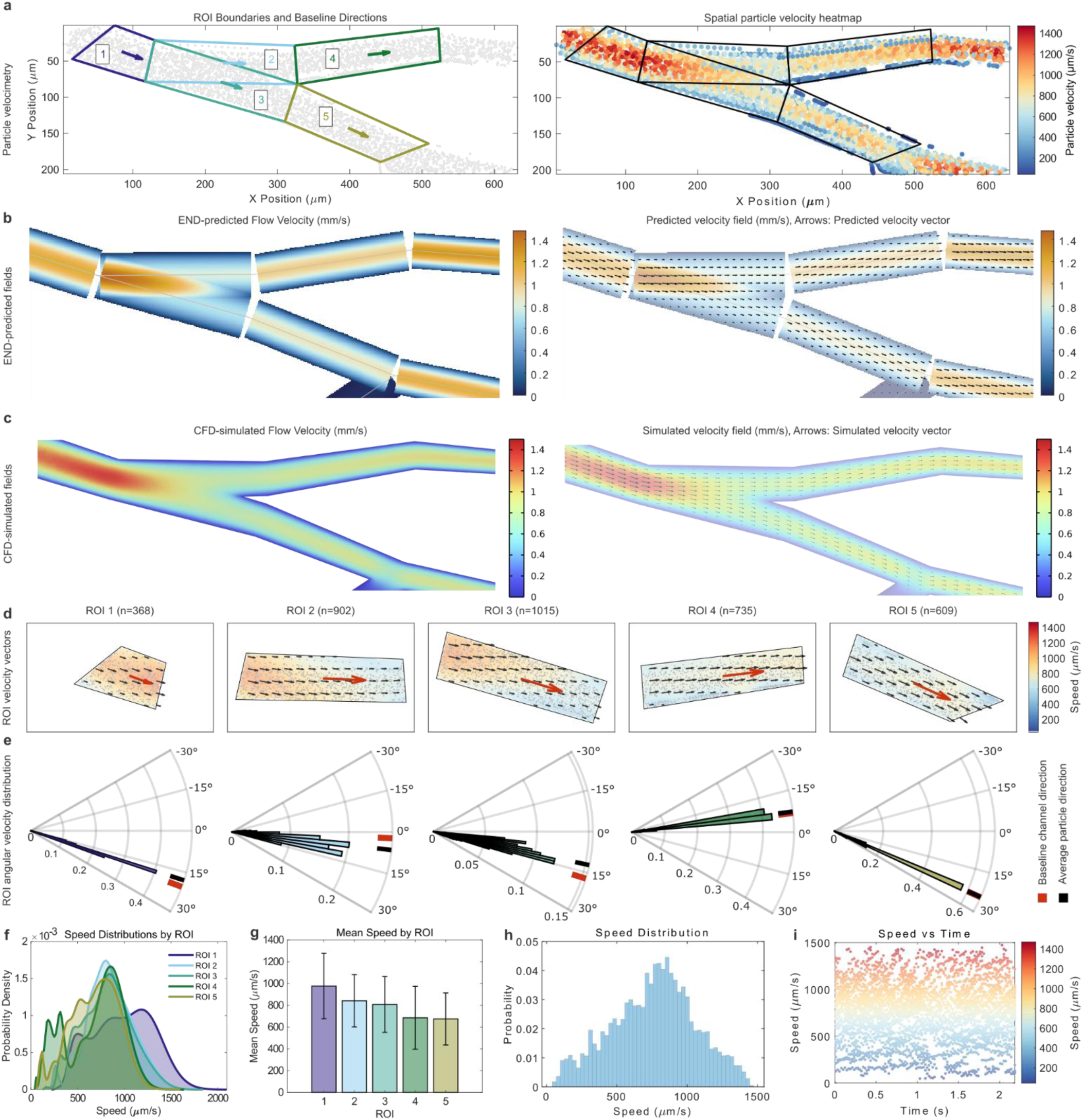
Spatial particle velocimetry reveals region-specific flow dynamics and cross-validates computational predictions. (**a**) Spatial analysis framework: Left panel shows Y-junction microfluidic device with five defined regions of interest (ROI 1-5, colored boundaries) and baseline channel direction indicators (arrows). Gray points represent individual particle trajectories captured via microscopy. Right panel displays spatial particle velocity heatmap. (**b**) END predictions: Left panel shows flow velocity magnitude distribution computed from electrical analog model, demonstrating rapid calculation of network-scale characteristics. Right panel presents predicted velocity vector field (arrows). (**c**) CFD simulation: detailed velocity magnitude (left) and vector field (right). CFD captures subtle hemodynamic features and local dynamics not resolved by END but requires orders of magnitude more computational time. (**d**) Region-specific experimental velocity analysis. (**e**) Angular velocity distribution analysis: Polar plots quantifying flow directionality in each ROI relative to baseline channel direction. ROI 1 shows narrow angular distributions confirming aligned laminar flow. ROI 2 and 3 exhibits wider angular spread indicating flow bifurcation and directional divergence. ROI 4 and 5 return to narrow distributions aligned with respective branch orientations, confirming flow stabilization downstream of bifurcation. (**f**) Speed probability distributions by ROI. (**g**) Quantitative regional flow comparison: Mean particle speeds with standard deviations across ROIs. Statistical analysis confirms highest velocities in pre-bifurcation region (ROI 1), maintained through bifurcation throat (ROI 2, 3), followed by deceleration in outlet branches (ROI 4, 5). (**h**) Global velocity distribution: Histogram of all measured particle speeds (n=3835 trajectories, including ones not ROIs). (**i**) Temporal velocity dynamics: Scatter plot of individual particle trajectory speeds versus time demonstrates stable flow conditions throughout 2-second acquisition period. Color gradient indicates speed magnitude. All color bars are set to have the same bounding range for better colormap comparison.

### Simulation of in vivo hemodynamics with whole blood

To extend validation beyond Newtonian fluid approximations and demonstrate physiological relevance, we perfused devices with simulated whole blood, simulating the multiphase transport characteristics of in vivo microcirculation. Red blood cell (RBC) motion was quantified using high-speed confocal microscopy, and cell speeds were extracted for individual segments (Fig. 10).

**Fig. 10:**
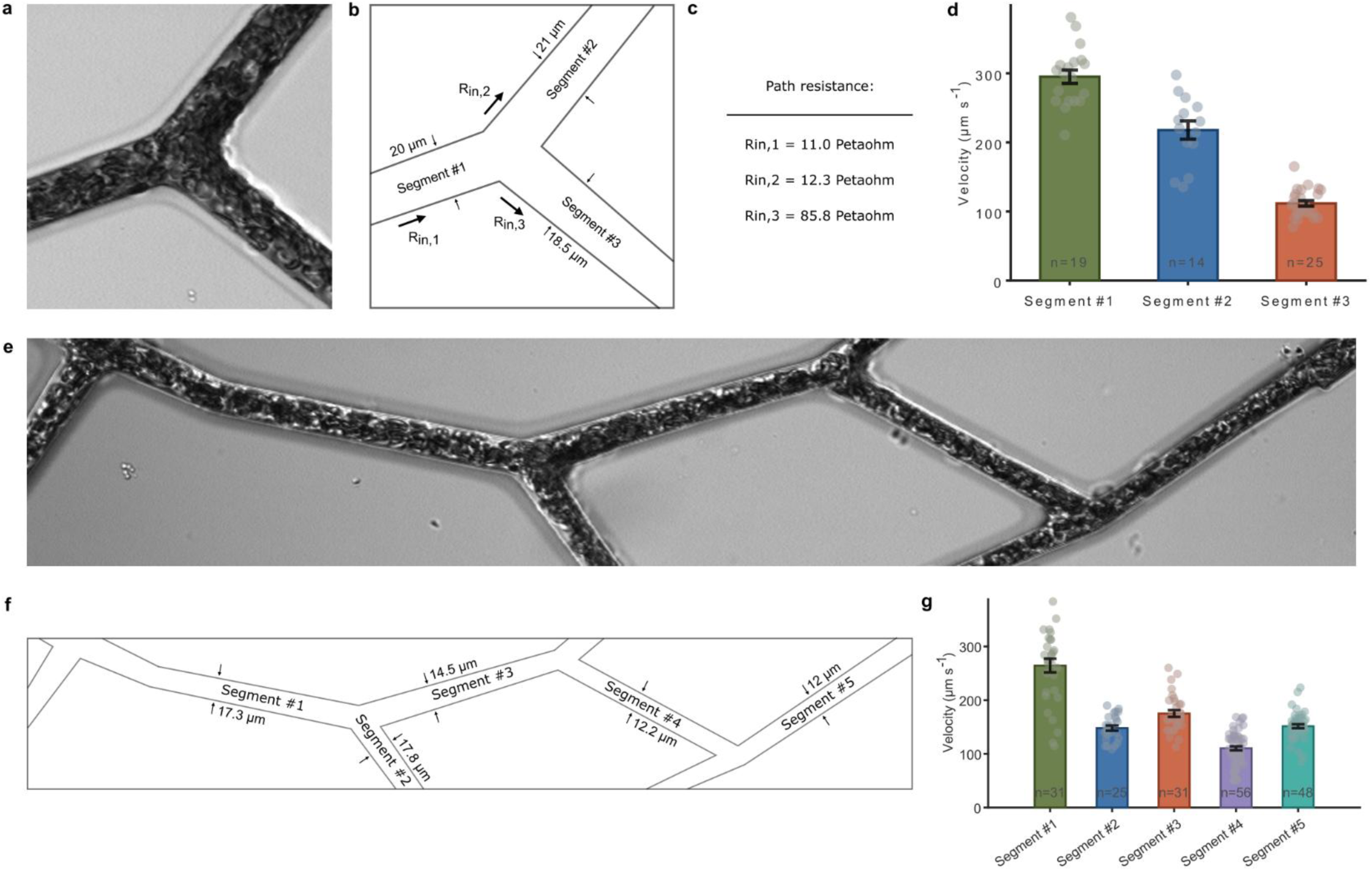
Simulation of in vivo hemodynamics with simulated blood in biomimetic microvascular networks. (**a**) High-speed confocal microscopy of red blood cell (RBC) transit through a single bifurcation perfused with simulated whole blood. (**b**) Schematic of the bifurcation geometry with segment labels and measured channel dimensions. (**c**) END-computed path resistances for each segment. (**d**) RBC speeds are extracted via cell tracking. (**e**) Confocal imaging of RBC flow through a multi-segment microvascular network. (**f**) Schematic of the network geometry with segment labels. (**g**) RBC speeds are extracted via cell tracking.

Cell tracking analysis revealed that RBC speeds and distribution patterns closely mirrored the heterogeneity predicted by our biologically informed network models. Segments identified computationally as high-resistance or low-resistance pathways exhibited correspondingly reduced or elevated RBC speeds, despite almost identical local geometries in many cases. The successful transit of blood cells through the varying diameters, ranging from larger arteriole-like channels down to capillary-scale (< 8µm) segments also confirmed that our inverse design algorithms effectively prevent the formation of unphysiologically narrow constrictions or stagnation zones that often lead to occlusion in synthetic networks. These results establish our platform’s capability to generate microfluidic environments that not only mimic the static structure of native vasculature but also recapitulate its dynamic functional behavior under complex, non-Newtonian conditions relevant to in vivo microcirculation.

## Discussion

The recapitulation of functional microvasculature remains a primary bottleneck in the advancement of organ-on-chip systems and tissue engineering. Current approaches have largely bifurcated into two domains: high-fidelity imaging of native tissue, which is difficult to manufacture, and reductionist microfluidic designs that sacrifice topological relevance for manufacturability(*41*). Our results establish a generative platform that integrates design, analysis, and fabrication of microvascular networks that captures defining architectural and functional features of biological capillary beds while remaining computationally and experimentally accessible. By unifying new approaches to stochastic, closed-circuit network generation, inverse parameter assignment, high throughput network-level characterization, and fabrication-aware translation within a single platform, our work addresses a long-standing gap between the complexity of native microvasculature and the simplicity of extant microfluidic vascular models.

A critical innovation presented here is the resolution of the “closed-circuit dilemma” regarding Murray’s Law. While Murray’s principle of minimum work is easily applied to branching tree designs(*40*), its application to closed-circuit networks has historically been computationally intractable due to circular dependencies in flow distribution(*36*). By exploiting the isomorphism between hydraulic and electrical circuits, our inverse design framework successfully decouples topology from geometry. This allows, for the first time, the automated generation of fabrication-ready anastomotic networks that rigorously satisfy biological scaling laws at every junction across scales and networks. This capability is essential for generating physiologically relevant shear stress gradients, which are known to drive endothelial phenotype and vascular remodeling(*4*, *3*). While recent advancements have enabled organ-scale vascular network generation and simulation, lack of compliance with Murray’s Law in closed loop design and fabrication limitations (∼400-μm minimum feature size) constrained the physiologic continuum of vessel sizes that could be modeled(*37*).

The introduction of the END framework represents a conceptual shift in how microvascular transport is modeled. By formalizing the isomorphism between the Hagen-Poiseuille equation and Ohm’s Law, we effectively reframe the problem from one of fluid mechanics to one of circuit theory. This is not merely a computational convenience but a methodological bridge that allows vascular engineering to leverage over 50 years of optimization in electronic circuit simulation. This approach decouples the calculation of network-level flux from the resolution of local flow fields. Traditional methods often conflate these scales, requiring millions of mesh elements to resolve a pressure drop that END determines analytically in milliseconds. Consequently, the END framework enables the rapid screening of thousands of architectures to identify functional outliers before investing computational resources in high-fidelity multiphysics simulations.

A central insight emerging from this work is that microvascular flow distribution is an emergent property of global network topology rather than local geometry(*42*). Our results demonstrate that in closed-circuit architectures, segments with identical dimensions can experience vastly different fluid dynamic environments depending solely on their connectivity within the larger network. This finding challenges the utility of simplified microfluidic models that assume local geometry dictates shear stress, suggesting that commonly used bifurcating trees may fundamentally misrepresent the mechanical cues present in biological capillary beds. The quantitative agreement between our computational predictions and particle velocimetry data confirms that our END framework accurately captures these topology-driven flows.

Together, our experiments establish a direct, quantitative correspondence between designed network architecture, computational prediction, and empirical measurement across modalities and perfusates. Identical microvascular networks produced consistent flow hierarchies in silico and in empirical devices at multiple experimental scales, creating true “digital twins” for vascular engineering. Our cross-platform validation confirms that the framework captures fundamental, topology-driven determinants of microvascular flow. By demonstrating predictive agreement from abstract network design through fabrication and biological perfusion, these results position our platform as a true “digital twins” for vascular engineering. Collectively, these tools have the potential to transform microvascular design from a trial-and-error process into a deterministic engineering discipline, providing a robust testbed for interrogating structure-function relationships in hematology, oncology, and regenerative medicine.

## Materials and Methods

### Network graph generation algorithm

We developed a stochastic growth-based algorithm in MATLAB to generate biology-informed microvascular networks within any given square domain size. The algorithm initializes with an arterial entry point at the left edge midpoint and an outlet at the right edge midpoint, creating initial vessel sprouts that advance through iterative growth steps of the given step size. Vessel growth direction is determined by combining parental momentum, branch memory, stochastic angular perturbations, and position-dependent outlet attraction. The domain is functionally divided into three regions: an early exploratory region with high directional randomness, a middle region where branching probability increases to maximize capillary density with wider angular variation, and a late region where vessels converge toward the outlet with reduced branching and narrower angles. Branching occurs stochastically with a given base probability, modulated by position, with minimum intervals of given number of nodes between successive branches along the same vessel to prevent local clustering. Branch angles incorporate the parent vessel direction weighted by memory strength, random offsets scaled by position, and small jitter components for natural irregularity. Closed-loop architecture emerges through two anastomosis mechanisms: forced connections when growing tips intersect existing segments, and proximity-based connections when tips approach within a given threshold of nodes or segment midpoints, with loop size verification preventing unrealistically small loops. All the model variables could follow a function instead of being constant. Parabolic function-assigned vessel diameters follow a parabolic profile with maximum diameter at domain edges and minimum at the center. Growth terminates when vessels exit domain boundaries, connect to the outlet, form anastomoses, or reach iterations threshold. Post-processing connects residual dangling tips in the right half to the outlet and integrates left-side danglers to nearby well-connected nodes (degree > 1) within a certain threshold. The algorithm generates networks with stochastic variation controlled by random seed selection enabling diverse network library generation from identical parameters.

### Inverse design of Murray’s Law-compliant diameters through electrical simulation

Microvascular networks exhibit diameter distributions governed by Murray’s Law, which relates vessel diameters to flow rates through power-law relationships. To computationally assign biologically realistic diameters to generated network topologies, we exploited the mathematical equivalence between fluidic networks obeying Murray’s Law and electrical circuits governed by Kirchhoff’s laws.

Each microvascular network topology (nodes and segments) from our computational library was mapped to an equivalent electrical circuit component. Vessel segments were represented as 1 Ω resistors connecting their respective endpoint nodes, with each node assigned a unique numerical identifier. Boundary conditions were implemented as a 1 V source between the terminal inlet node and ground. SPICE netlists were automatically generated using custom MATLAB scripts and solved via LTspice (Analog Devices) to determine steady-state branch currents (analogous to volumetric flow rates) through operating point analysis. The calculated branch currents were transformed into vessel diameters using Murray’s Law exponent: d ∝ I^(1/3), where d represents diameter and I represents electrical current. Raw diameter values were normalized by dividing by the minimum non-zero value, then mean-centered. The final diameter distribution was linearly scaled such that the minimum diameter corresponded to the desired scale. The assigned diameter data were stored in the model library as .mat files for subsequent fabrication and validation steps.

### 3D object creation for microfabrication

We developed a custom MATLAB-to-OpenSCAD pipeline to translate microvascular network graphs into 3D objects suitable for microfabrication. For each segment, we calculated geometric properties including length, midpoint coordinates, and orientation angle with negation to align with OpenSCAD’s coordinate system. Each node is assigned a diameter equal to the largest diameter of any connected segment to ensure smooth transitions at vessel junctions. The vascular network is geometrically represented as a union of truncated cones and spheres, where each segment was modeled as a cylinder with end diameters (d1, d2) matching its connected node diameters, positioned at the segment midpoint (translated relative to domain center), rotated to match segment orientation using a compound rotation [angle, 90, 0], with the cylinder length matching the calculated segment length and centered at its position. Spheres were placed at each node with diameters matching the node’s largest connected segment, positioned relative to the domain center. A vertical offset parameter was applied at an appropriate height. The entire vascular geometry was positioned within a Boolean difference operation that subtracted a large cube to create a flat base plane and remove geometry below z = 0. For fabrication purposes, inlet and outlet port assemblies were added at opposing domain edges. The MATLAB code generated OpenSCAD scripts by constructing string arrays beginning with rendering quality parameters followed by Boolean operations (difference and union), geometric primitives (cylinder, sphere, cube) with transformation commands (translate, rotate), and numerical parameters derived from network data, which were exported as text files and subsequently executed in OpenSCAD software to generate STL files for fabrication processes.

### 3D Microfabrication

The rendered 3D models were fabricated using a Nanoscribe GT Photonics Professional two-photon polymerization (2PP) 3D printing system. Four-inch silicon wafers were used as printing substrates. Wafers were sequentially washed with acetone and isopropanol (IPA) to remove organic contaminants, rinsed with deionized (DI) water, dried under nitrogen flow and dehydrated on a hotplate at 150°C for 5 minutes to enhance resin adhesion. The structures were printed using IPX-Q resin (NanoScribe). After printing, the resin was developed for 20 minutes in SU-8 developer followed by an IPA rinse. The wafer was post-cured in a UV chamber (Formlabs cure) for 20 minutes to cure the uncured resin. A layer of (Tridecafluoro-1,1,2,2-Tetrahydrooctyl)Trichlorosilane was applied to the wafer. PDMS replicas (10:1 ratio) of the master mold were made and bonded to glass slides using oxygen plasma to form the final devices. Bonded devices underwent quality control inspection using optical microscopy to verify channel continuity and seal integrity.

### 2D mask creation for microfabrication

We developed a MATLAB pipeline to generate photomask designs for soft lithography fabrication by converting microvascular network graphs into GDS II format using the GDS II toolbox (*43*) library. Each node was assigned a diameter equal to the largest diameter of any connected segment to ensure complete coverage at vessel junctions. The network geometry was represented as a combination of paths and filled circles, where vessel segments were drawn as paths with widths matching segment diameters, connecting node coordinates, and filled circles were placed at each node with diameter equal to the node’s maximum diameter to eliminate gaps at segment intersections caused by angular connections. Inlet and outlet port structures were added by identifying nodes at domain boundaries (maximum and minimum x-coordinates), extending horizontal paths from these nodes with widths matching the boundary node diameters, and terminating each path with a circular reservoir (1 mm diameter) to facilitate tube connections in the final microfluidic device.

### 2D microfabrication

2D PDMS-based microfluidic devices were fabricated using standard soft lithography techniques with master molds created via maskless photolithography. Silicon wafers were washed with Acetone and IPA, rinsed with DI water, dehydrated on a hotplate at 150°C for 10 minutes, and spin-coated with SU-8 negative photoresist following the manufacturer’s recommended processing parameters. The GDS II mask design was directly written onto the SU-8 photoresist using a Heidelberg Instruments MLA 150 maskless lithography system. Following exposure, wafers were processed according to standard SU-8 datasheet protocols. A layer of (Tridecafluoro-1,1,2,2-Tetrahydrooctyl)Trichlorosilane was applied to the wafer. PDMS replicas of the master mold were made and bonded to glass slides using oxygen plasma to form the final devices.

### 3D object creation for fluid dynamics simulations

We developed a custom MATLAB-to-COMSOL pipeline to translate microvascular network graphs into 3D geometries suitable for computational fluid dynamics simulations. For each segment, we calculated geometric properties including length (Euclidean distance between endpoints), endpoint coordinates (p1, p2), midpoint coordinates, and orientation angles using arctangent functions to determine both the azimuthal angle and the x- and y-components of the segment direction vector. Each node was assigned a diameter equal to the largest diameter of any connected segment to ensure smooth transitions at vessel junctions. The vascular network was geometrically represented as a union of truncated cones and spheres using COMSOL’s API commands. Each segment was modeled as a cone with base and top radii set to half the diameters of its connected nodes, height matching the segment length, positioned at the first endpoint (p1), and oriented using spherical coordinates with axis components derived from the segment direction vector. Spheres were placed at each node with radii set to 50.001% of the node’s largest connected segment diameter to ensure proper junction connectivity. The MATLAB code generated COMSOL Java API scripts as string arrays containing geometry creation commands, and numerical parameters derived from network data, which were exported as text files for execution in COMSOL Multiphysics to create computational geometry for finite element mesh generation and fluid dynamics simulations.

### Computational Fluid Dynamics

Computational fluid dynamics simulations were performed using COMSOL Multiphysics software with automated geometry generation via the COMSOL Java API. The 3D vascular geometries were created programmatically by executing the MATLAB-generated COMSOL API scripts, which constructed the network as a union of truncated cones and spheres as described previously, ensuring direct correspondence between the computational domain and the experimentally fabricated structures. The fluid domain representing the vessel lumens was assigned water as the working fluid (density = 1000 kg/m³, dynamic viscosity = 0.001 Pa·s for water) to enable initial model validation and comparison with microfluidic experiments, though blood rheology models can be implemented for more physiologically accurate simulations. Steady-state laminar flow simulations were conducted using COMSOL’s finite element method, with appropriate boundary conditions applied at the inlet and outlet ports (specified pressure differential or flow rate at inlet, zero pressure at outlet) and no-slip conditions at vessel walls. The computational mesh was generated using COMSOL’s automatic meshing algorithms. Simulation outputs included spatial distributions of pressure profiles throughout the vascular network and velocity vector fields and magnitude profiles showing flow patterns and preferential flow pathways.

### Electrical network dynamics (END)

To rapidly estimate pressure and flow distributions in the microvascular networks, we employed an electrical circuit analog model based on the Hagen-Poiseuille equation, where hydraulic resistance, pressure, and volumetric flow rate are analogous to electrical resistance, voltage, and current, respectively. Each vessel segment was modeled as a fluidic resistor with hydraulic resistance calculated using a modified Hagen-Poiseuille equation accounting for specific channel cross-sectional geometries. For channels with irregular shapes, a geometric correction factor (0.8538 for 75% circle) accounts for the reduced cross-sectional area compared to full circular channels. Custom MATLAB scripts automatically generate SPICE (Simulation Program with Integrated Circuit Emphasis) netlist files representing each microvascular network as an electrical circuit. Each node was assigned a unique numerical identifier (0 to N-1), vessel segments were represented as resistors connecting their endpoint nodes with resistance values derived from the Hagen-Poiseuille equation, and boundary conditions were implemented as a 1 V DC source connected between the terminal inlet node and ground, representing a 1 Pa inlet-outlet pressure differential. The generated SPICE netlists were solved using LTspice (Analog Devices) with DC operating point analysis (.op command) to determine steady-state nodal voltages (analogous to pressures) and branch currents (analogous to volumetric flow rates). The total circuit current (I_circuit_) was extracted from the voltage source element, and the equivalent network hydraulic resistance was calculated as R_eq_ = 1/(I_circuit_). Nodal voltages and branch currents were parsed from LTspice “.raw” output files using LTSPICE2MATLAB function (*44*) for subsequent flow analysis and visualization.

### Model resistance calculations

In END, the equivalent network resistance was calculated from SPICE simulation results by: R = V_source_ / I_circuit_, where V_source_ is the applied voltage (1 V, analogous to 1 Pa pressure differential) and I_circuit_ is the total current through the voltage source (analogous to total volumetric flow rate).

For CFD simulations, network resistance was computed from simulation outputs as R = ΔP / Q, where ΔP is the pressure difference between inlet and outlet (0 Pa) and Q is the volumetric flow rate. Simulations were performed with outlet pressure set to zero (atmospheric reference), a specified inlet flow rate, and the resulting inlet pressure measured post-simulation to determine the pressure drop across the network.

For fabricated in vitro microfluidic devices, hydraulic resistance was measured under controlled flow conditions. Devices were connected to a syringe pump operating at a specified volumetric flow rate, with the outlet channel at atmospheric pressure (P_out_ = 0). Inlet pressure (P_in_) was monitored in real-time using an in-line pressure sensor (EIPS345, Fluigent Inc.). Network resistance was calculated as R = P_in_ / Q.

### Particle tracing and flow field profiles

Fluorescent Pink Particles (Spherotech, Lake Forest, IL) with a size of 0.92 micron and 1:200 PBS dilution of original 1.0% w/v concentration were used to conduct particle tracing experiments. Series images were captured using a 20X objective with a FLUOVIEW FV4000 Confocal Laser Scanning Microscope. The images were processed using TrackMate within FIJI (ImageJ). Sequential frames were used to determine the flow directions with each particle’s movement translated into a velocity field vector. Track data was exported for analysis.

### Software and analysis

We used MATLAB R2023a (MathWorks) for creating the platform and the pipelines; MATLAB R2023a (MathWorks) or R(RStudio) to analyze the data and create raw plots; Excel (Microsoft), COMSOL 6.3 (COMSOL AB), or MATLAB R2023a (MathWorks) to record data; LTspice 24.1.9 (Analog Devices) for electrical simulations; KLayout (klayout.de) for GDS inspections; FIJI (ImageJ) or Inkscape (inkscape.org) for image analysis and figure generation; Trackmate(*45*) for object tracing; OxyGEN (FLUIGENT) for real-time pressure readings; and OpenSCAD 2021.01 (openscad.org) for 3D object generation.

### Blood experiments

To extend validation beyond Newtonian fluid approximations and demonstrate physiological relevance, we perfused selected devices with simulated whole blood to evaluate the multiphase transport characteristics of in vivo microcirculation. Whole blood was collected in ETDA tubes under approved IRB protocol. Red blood cell (RBC) fraction was isolated using Ficoll density gradient separation (1.077g/mL) motion was quantified using high-speed confocal microscopy, and cell speeds were extracted for individual segments and junctions.

## Acknowledgements

The authors would like to thank the staff at Georgia Tech Institute for Matter and Systems for their support and assistance in the fabrication and characterization of the microdevices; the McCallum Early Career Award (S.N.H.), grants from the Northside Hospital Foundation, Inc. (S.N.H.), the Ovarian Cancer Institute (S.N.H.), and Georgia Tech IMS (A.V. and S.N.H.) for their support. This work was performed in part at the Georgia Tech Institute for Matter and Systems, a member of the National Nanotechnology Coordinated Infrastructure (NNCI), which is supported by the National Science Foundation (Grant ECCS-2025462). Colorblind-safe color schemes from Paul Tol’s technical note (https://sronpersonalpages.nl/~pault/data/colourschemes.pdf) were used in visualizations of this work.

## Author contributions

A.V. and S.N.H. conceived this project, designed the microvascular generation logic, and wrote the initial manuscript. A.V. designed the END pipeline, Inverse Current-to-Diameter Transformation (ICDT), and hybrid END-CFD approach, wrote the software, fabricated the in vitro devices, and generated visualizations. S.N.H. supervised the study and provided resources and funding. A.V., A.R.B., and S.N.H. conducted the experiments, analyzed the data, reviewed, and edited the manuscript before submission.

## Competing interests

A.V. and S.N.H. are named inventors on a provisional patent application filed by Georgia Tech that covers the technology related to this work.

## Data availability

The data that support the findings of this study are available on request from the corresponding author.

